# Cytosolic transport of citrate protects nutrient-austere pancreatic cancer from ferroptosis

**DOI:** 10.1101/2025.01.15.633097

**Authors:** Adam Kneebone, Kailey Lindaur, Ata Abbas, Joel Cassel, Sarah Graff, Gerard Abood, Xianzhong Ding, William Small, Curtis Tatsuoka, Simone Sidoli, Joseph M. Salvino, Ali Vaziri-Gohar

## Abstract

Pancreatic cancer (PDAC) cells experience nutrient starvation in a poorly perfused tumor microenvironment. Metabolic dependencies that protect PDAC cells from detrimental oxidative stress in a nutrient-restricted niche represent as tumor-specific targets. While the role of mitochondria in supporting energy production and biosynthetic requirements of cells has been well investigated, their contribution to maintaining intracellular redox homeostasis when PDAC cells are exposed to nutrient deprivation is unknown. Our results demonstrate that cytosolic transport of citrate via SLC25A1 confers a survival advantage to PDAC cells by protecting them from ferroptosis, a well-established iron-dependent cell death mechanism, under nutrient-limited conditions. Employing selective SLC25A1 inhibitor or targeting mitochondrial OXPHOS dramatically reduced GPX4 expression and PDAC cell viability. Rescuing GPX4 expression with the products of both ACLY and ACO1-dependent pathways uncovered their critical role in conferring survival advantage under metabolic stress. Importantly, exogenous expression of GPX4 reversed redox imbalance and metabolic discordance resulting from the lack of SLC25A1 activity, indicating the requirement of citrate-induced GPX4 expression to support mitochondrial health and function. As observed with cultured cells under nutrient limitation, SLC25A1 function was revealed to be indispensable in pancreatic tumor microenvironment, and the reduced growth, due to the lack of SLC25A1 activity, was rescued with antioxidant NAC in preclinical models of PDAC. Lastly, SLC25A1 suppression was accompanied by elevated glutamine metabolism, and combination therapy with pharmacologic inhibitors of SLC25A1 and glutaminase inhibitor CB-839 dramatically suppressed tumor growth, highlighting this combinatorial approach as a potential therapeutic strategy in PDAC.

## Main

Pancreatic cancer (PDAC) cells live in a microenvironment characterized by reduced elemental nutrients, such as glucose, resulting from insufficient vascularization^1–4^. Nutrient limitation generates oxidative stress which can limit cancer growth^1,5^. To maintain growth in nutrient-austere environment, PDAC cells recruit mechanisms that support generation of reducing potential, such as metabolic activities generating NADPH and antioxidant glutathione, to eliminate excess reactive oxygen species (ROS)^1,6–8^. Emerging evidence indicates mitochondrial activities are crucial to supporting PDAC survival, particularly when nutrient availability is limited^1,9–11^. Mitochondrial biogenesis and oxidative phosphorylation (OXPHOS) are upregulated in cells under glucose limitation^1,6,10,11^. Mitochondrial ability to support cellular bioenergetic and biosynthetic requirements has been well investigated^12,13^; however, their potential role in terms of maintaining intracellular redox balance under nutrient limitation is unknown. We sought to determine whether mitochondrial activity is functionally linked with redox homeostasis when PDAC cells experience nutrient deprivation, the hallmark of pancreatic tumor microenvironment.

### Citrate supplementation rescued cells from oxidative stress results from ETC suppression

To determine if mitochondria protect cells from oxidative stress under nutrient limitation, PDAC cells were treated with sublethal doses of mitochondrial ETC inhibitors rotenone and antimycin A, the inhibitors of complex I and complex III, respectively, to block mitochondrial respiration. Upon OXPHOS inhibition, an increase in mitochondrial-derived ROS and total intracellular ROS levels were characterized by MitoSOX and DCFDA assays, respectively (**Fig. 1a, b**), suggesting mitochondrial activities impact redox balance globally when PDAC cells experience nutrient stress. Interestingly, oxidative insults were partially rescued within mitochondria by adding a mitochondrial-targeted antioxidant (Mito-Tempo); however, the treatment did not rescue the entire cell from excess ROS (**Fig. 1a, b**). These results suggest mitochondria play a substantial role, beyond its own compartment, in maintaining intracellular redox homeostasis.

**Figure 1.**
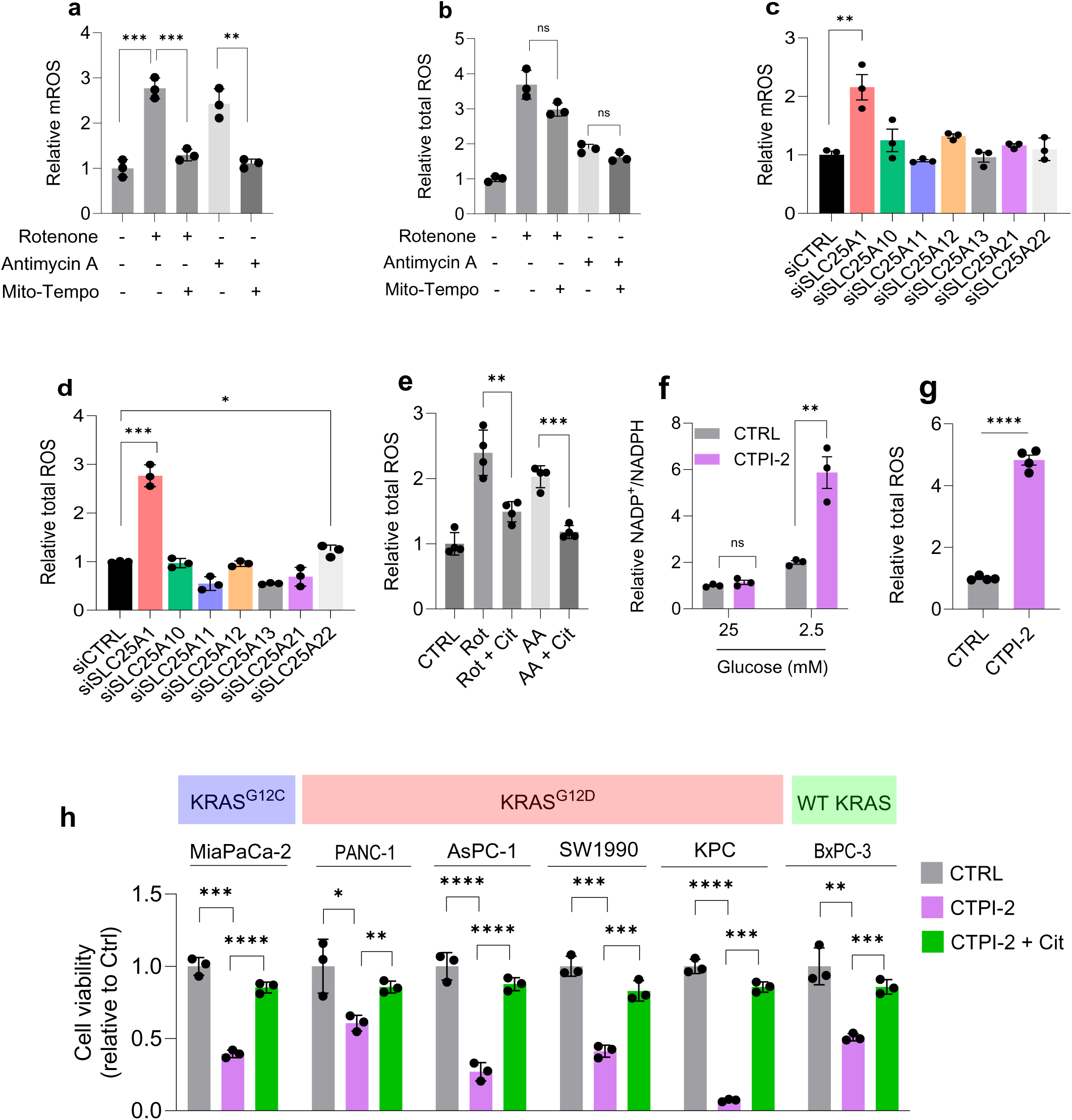
SLC25A1 is required for maintaining redox balance in PDAC cells under nutrient stress. **a, b,** Relative mitochondrial-derived ROS and total intracellular ROS detected by MitoSOX and DCFDA assays, respectively, in MiaPaCa-2 cells treated with Rotenone (100 nM), Antimycin A (100 nM), and Mito-Tempo (100 µM) for 48 hours. **c, d,** Relative mitochondrial-derived ROS and intracellular ROS in cells were incubated with siRNAs against indicated mitochondrial transporters for 72 hours followed by incubation under low glucose conditions for an additional 48 hours. **e,** Relative intracellular ROS levels in cells treated with Rotenone (Rot, 100 nM), Antimycin A (AA, 100 nM), cotreatment with sodium citrate (Cit, 4mM), or control for 48 hours. **f,** Relative NADP+/NADPH ratio in cells treated with selective SLC25A1 inhibitor CTPI-2 (100 µM) or control under indicated conditions for 48 hours. **g,** Relative total ROS levels in cells treated with CTPI-2 (100 µM) or control under glucose limitation for 72 hours. **h,** Relative viability of various PDAC cells treated with CTPI-2 (100 µM), cotreatment of CTPI-2 and sodium citrate (Cit, 4mM), or control for 5-7 days. n=3 independent biological replicates, unless indicated. Data provided as mean±s.d. Pairwise comparisons were conducted using two-tailed, unpaired Student’s *t*-tests. *representing *p*<0.05,**representing *p*<0.01, ***representing *p*<0.001, ****representing *p*<0.0001.

To explore this, we performed an unbiased siRNA screen of all mitochondrial transporters enabling transport of TCA cycle intermediates to the cytosol. Among these transporters, silencing of SLC25A1 exhibited a robust increase in both mitochondrial-derived and intracellular ROS in cells cultured in glucose-limited medium (**Fig. 1c, d**). SLC25A1, aka citrate transport protein, mediates the transport of mitochondrial-derived citrate to the cytosol^14^. Further investigation showed that SLC25A1 mRNA levels were upregulated in PDAC cell lines under low glucose conditions, compared to the control which cultured in glucose abundance (**Extended Data Fig. 1a**). We also assessed whether citrate supplementation can rescue cells from observed oxidative insults derived from ETC inhibition.

Extracellular citrate can enter the cells via plasma-membrane localized citrate transporter SLC13A5^15^. Metabolic tracing study can be employed to monitor the entrance of exogenous citrate to the cells. Using [1,5,6-^13^C_3_]citrate, we showed the enrichment of citrate (M+3) in the cells (**Extended Data Fig. 1b**). Citrate supplementation normalized ROS levels to the basal level in cells treated with ETC inhibitors (**Fig. 1e**), indicating that mitochondria contribute to cellular redox balance, in part, by providing citrate to the cytosol. To further validate these findings, PDAC cells were treated with pharmacologic SLC25A1 inhibitor CTPI-2^16–18^. Cells experienced oxidative stress substantially under nutrient limitation (**Fig. 1f, g**), and CTPI-2 treatment led to a robust cell death, and this phenotype was reversed when cells were co-treated with citrate, as shown in a panel of PDAC cell lines (**Fig. 1h**), further validating on-target effects of the inhibitor. Taken together, these results revealed a role for SLC25A1 in supporting redox balance in PDAC cells under nutrient stress.

### SLC25A1 protects cells from ferroptosis under nutrient limitation

The underlying mechanism by which SLC25A1 inhibition impacts cell growth has not yet described. Transcriptomic studies revealed differentially expressed genes upon SLC25A1 knockdown relative to control (**Fig. 2a**). Among inhibitors of known cell death mechanisms, co-treatment of CTPI-2 with pharmacologic inhibitors of ferroptosis, including NAC (N-acetyl-cysteine), Fer-1, and Trolox, rescued cell viability in PDAC cells (**Fig. 2b**). Increased lipid peroxidation is a surrogate marker of ferroptosis-mediated cell death^19,20^. Our observations with employing various cell death inhibitors were corroborated by lipid peroxidation levels in cells under CTPI-2 treatment. Treating cells with CTPI-2 significantly induced lipid peroxidation in PDAC cells, suggesting ferroptosis as a potential consequence of inhibiting mitochondrial export of citrate (**Fig. 2c**).

**Figure 2.**
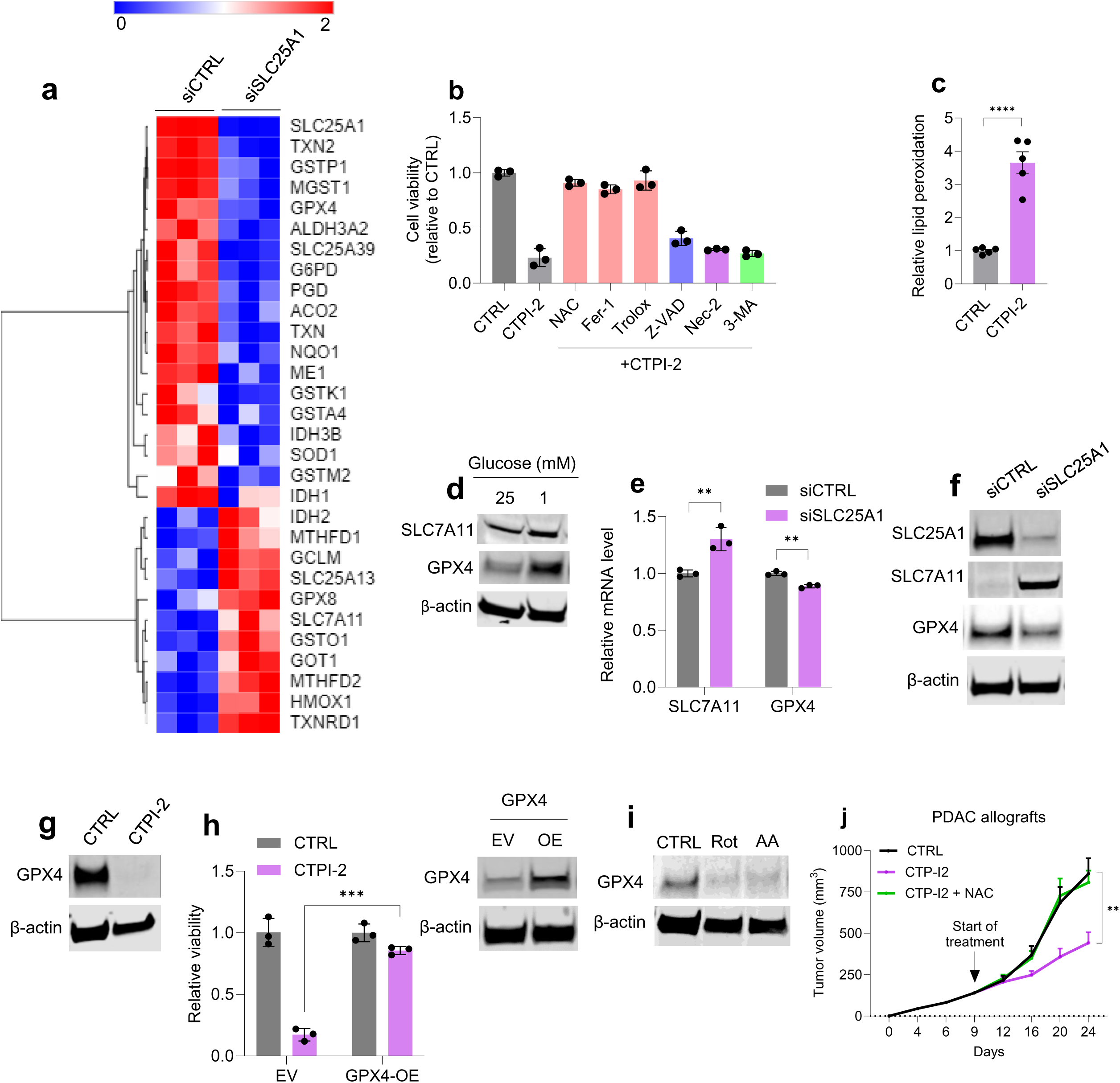
SLC25A1 protects PDAC cells from ferroptosis via regulation of GPX4 expression under nutrient limitation. **a,** Heatmap of differentially expressed genes associated with antioxidant defense in cells incubated with siRNA against SLC25A1 compared to control siRNA. **b,** Relative survival of cells treated with CTPI-2 (100 μM) alone or in combination with NAC (N-acetylcysteine, 1 mM), Fer-1 (2 μM), Trolox (100 μM), Z-VAD (10 μM), Nec-2 (10 μM), or 3-MA (10 μM) for five days. **c,** Lipid peroxidation measurements in cells cultured in medium containing 1LmM glucose and treated with CTPI-2 (100LµM) or control for 72 hours. **d,** Immunoblots of SLC7A11 Immunoblot analysis to evaluate SLC7A11 and GPX4 protein expression in cells cultured in medium containing either 25 mM glucose or 1 mM glucose for 24 hours. **e,** Relative mRNA levels of indicated genes, as determined by RNA sequencing, in SLC25A1 knockdown cells relative to control. **f,** Immunoblots of SLC25A1, SLC7A11, GPX4, and b-actin (control) in indicated cells. **g,** Immunoblots of GPX4 and control in cells treated with CTPI-2 (100 uM) for 72 hours. **h,** Relative survival of cells transfected with either empty vector or GPX4 overexpressing vector and treated with indicated treatments in medium containing 1 mM glucose. Western blot validation of GPX4-OE in MiaPaCa-2 cells under glucose abundance. **i,** Immunoblots of GPX4 and control in cells treated with Rot (rotenone, 100 nM), AA (antimycin A, 100 nM), or control for 48 hours. **j,** The growth of subcutaneous KPC allograft tumors in a syngeneic mouse model of PDAC with indicated treatments. CTPI-2 (30 mg/kg, IP, once daily; NAC (1 g/L in drinking water). n=6 mice per group. n=3 independent biological replicates, unless indicated. Data provided as mean±s.d. Pairwise comparisons were conducted using two-tailed, unpaired Student’s *t*-tests. **representing *p*<0.01, ***representing *p*<0.001, ****representing *p*<0.0001.

Protection against ferroptosis occurs via sufficient production of antioxidant glutathione and enzymatic activity of GPX4 for detoxification of lipid ROS^19–21^. To gain mechanistic insights on how the lack of SLC25A1 activity induced ferroptosis in cells exposed to nutrient limitation, we determined expression of key components of the cellular mechanism against ferroptosis. We thus aimed to profile the expression of these elements in cells under nutrient deprivation compared with cells under nutrient abundance. Culturing cells in nutrient limitation did not change the expression of SLC7A11, the protein involved in cellular transportation of cystine, whereas it increased the expression of GPX4 (glutathione peroxidase 4) in cells (**Fig. 2d**). The mechanisms regulating expression of GPX4 under nutrient limitation are unknown. While silencing SLC25A1 expression with siRNA did not reduce expression of SLC7A11, a reduction in GPX4 mRNA and protein expression was detected (**Fig. 2e, f and Extended Data Fig. 2**), and similar results were observed with CTPI-2 treatment (**Fig. 2g**), further indicating that cells lacking SLC25A1 expression or activity suffer from oxidative stress due to induction of ferroptosis.

To further investigate whether the lack of GPX4 expression was the underlying mechanism of observed cell death in cells treated with CTPI-2, the expression of GPX4 was exogenously increased. Overexpression of GPX4 markedly restored viability of cells under treatment with the inhibitor (**Fig 2h**), providing proof that the lack of SLC25A1 activity induces ferroptosis, which then suppresses cell viability in nutrient-restricted PDAC cells. We subsequently assessed if mitochondria via SLC25A1 promote expression of GPX4 under glucose limitation. As expected, a robust reduction in GPX4 protein levels were observed upon OXPHOS inhibition in PDAC cells under low-glucose conditions (**Fig. 2i**), validating the role of mitochondria in protecting cells from ferroptosis under metabolic stress. Furthermore, these findings translated into a robust therapeutic approach in mice with allografts tumors. Treating with SLC25A1 inhibitor CTPI-2 suppressed tumor growth, and providing mice with NAC completely rescued the phenotype (**Fig. 2j**), further validating the role of SLC25A1 in supporting redox balance in PDAC tumors.

### Both ACLY and ACO1 mediated pathways regulates GPX4 expression

Cytosolic citrate, provided by SLC25A1, can supply substrate for two separate pathways: an upstream ACLY (ATP-citrate lyase)-dependent pathway that leads to Acetyl-CoA and fatty acid synthesis^22,23^ and a downstream aconitase 1 (ACO1)-dependent pathway which mediates the generation of various metabolites and reducing molecules, such as αKG, glutamate, and reducing agent NADPH, via a series of metabolic reactions^1,6,24^ (**Fig. 3a**). The SLC25A1 support of the downstream pathway mediates activation of numerous metabolic reactions, such as IDH1 and GOT1 which have been previously shown to play an essential role for pancreatic cancer growth and survival^1,25^. In order to determine the key metabolic pathway that is mediating the role of SLC25A1 in GPX4 expression, we first investigated whether cytosolic citrate serves as a substrate for the abovementioned pathways. Indeed, the products of these pathways, including acetyl-CoA, which can be produced from acetate, αKG, and glutamine (the precursor of glutamate) rescued GPX4 expression and restored cell viability in cells lacking SLC25A1 activity via gene knockdown or pharmacologic inhibition of the transporter (**Fig. 3b, c**). Acetate can generate acetyl-CoA via the activity of acetyl-CoA synthetases^26^, and ACSS2 expression was determined in the cytosolic fraction (**Extended Data Fig. 3a**). Consistent with these results, siRNA-mediated knockdown of enzymes linked with cytosolic citrate, ACLY and ACO1, markedly diminished GPX4 expression and caused upregulation of ROS levels (**Fig. 3d, e**), further establishing the contribution of these metabolic nodes to supporting PDAC survival under nutrient limitation.

**Figure 3.**
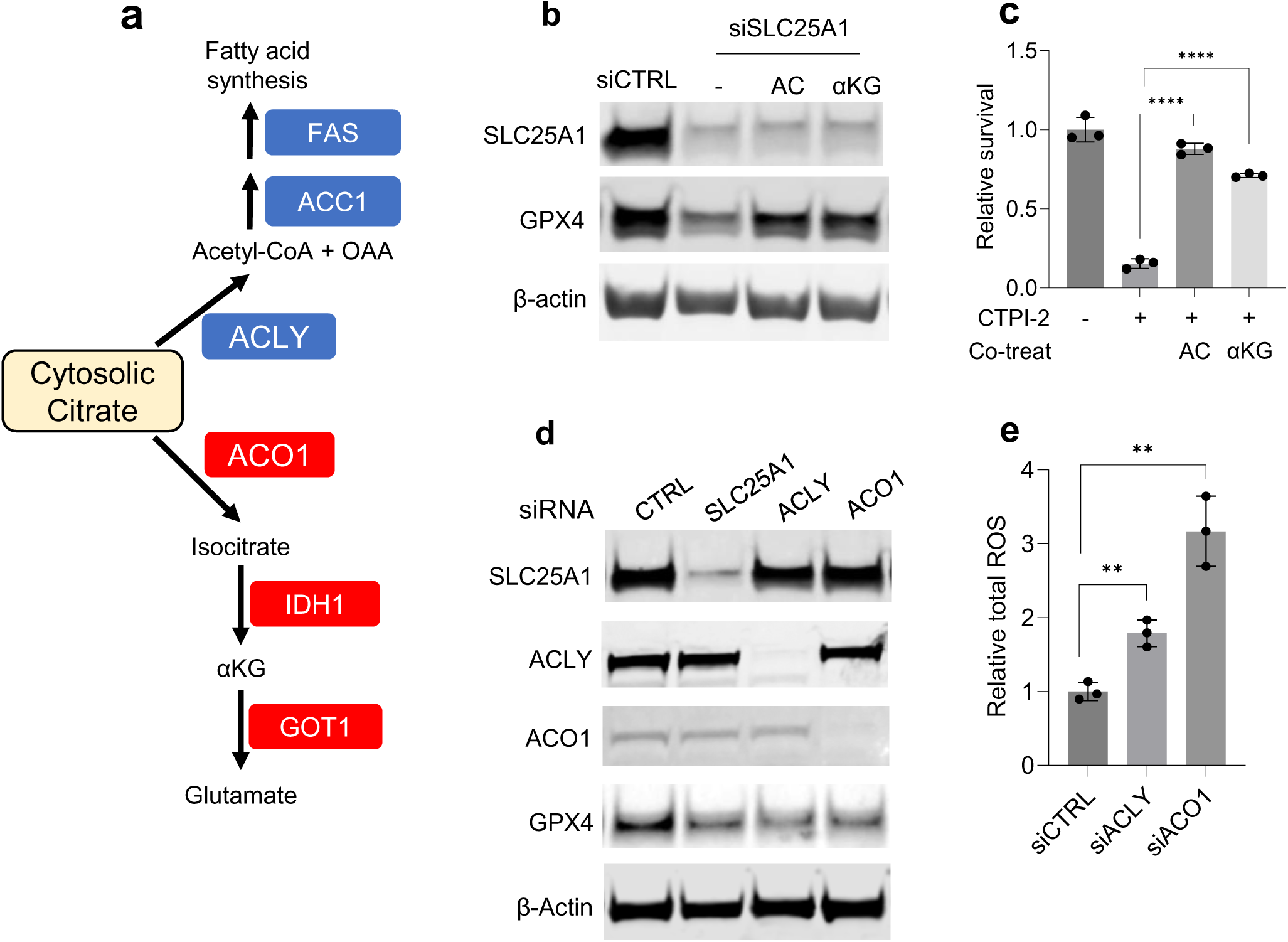
Cytosolic transport of citrate activates both ACLY- and ACO1-depenent reactions to promote GPX4 expression under nutrient-limited conditions. **a,** Schematic illustrating metabolic pathways supported by cytosolic citrate. ACLY, ATP-citrate lyase; OAA, oxaloacetic acid; ACO1, aconitase 1; IDH1, isocitrate dehydrogenase 1; aKG, alpha-ketoglutarate; GOT1, cytosolic aspartate aminotransaminase. **b,** Immunoblots of SLC25A1, GPX4, and control in SLC25A1 knockdown cells and control after incubation with sodium acetate (AC, 4mM) and α (4mM) for 48 hours. **c,** Relative survival of cells under indicated treatments for 5-7 days. CTPI-2 (100 µM). Immunoblots of SLC25A1, ACLY, ACO1, GPX4, and control (**d**) and relative total intracellular ROS levels (**e**) in indicated knockdown cells when incubated under glucose-limited (1 mM glucose) conditions for 48 hours. n=3 independent biological replicates). Data provided as mean±s.d. Pairwise comparisons were conducted using two-tailed, unpaired Student’s *t*-tests. **representing *p*<0.01, ****representing *p*<0.0001.

### SLC25A1 is required for histone acetylation under nutrient limitation

Acetyl-CoA produced via acetate can directly support histone acetylation. As shown, acetate supplementation led to a complete rescue of GPX4 expression and cell viability in cells under SLC25A1 inhibition (**Fig. 3b, c**), suggesting histone acetylation could play a critical role in SLC25A1-mediated GPX4 expression. While SLC25A1 knockdown mitigated total histone H3 acetylation, it did not alter histone H4 acetylation levels (**Fig. 4a-c**). Similar results were also reproduced by western blot and revealed histone H3 to be reduced in SLC25A1 knockdown cells compared to the control (**Fig. 4d**). The changes were especially pronounced at lysine 18 and 27 positions in the given cells under low glucose conditions (**Fig. 4e**). As observed with the rescue experiments of GPX4 expression with downstream metabolites (**Fig. 3b**). Incubating cells with acetate, αKG, and, glutamate precursor glutamine, exhibited similar effects on restoring histone H3-K27 acetylation in SLC25A1 knockdown cells (**Fig. 4f**), suggesting the contribution of histone acetylation to GPX4 induction.

**Figure 4.**
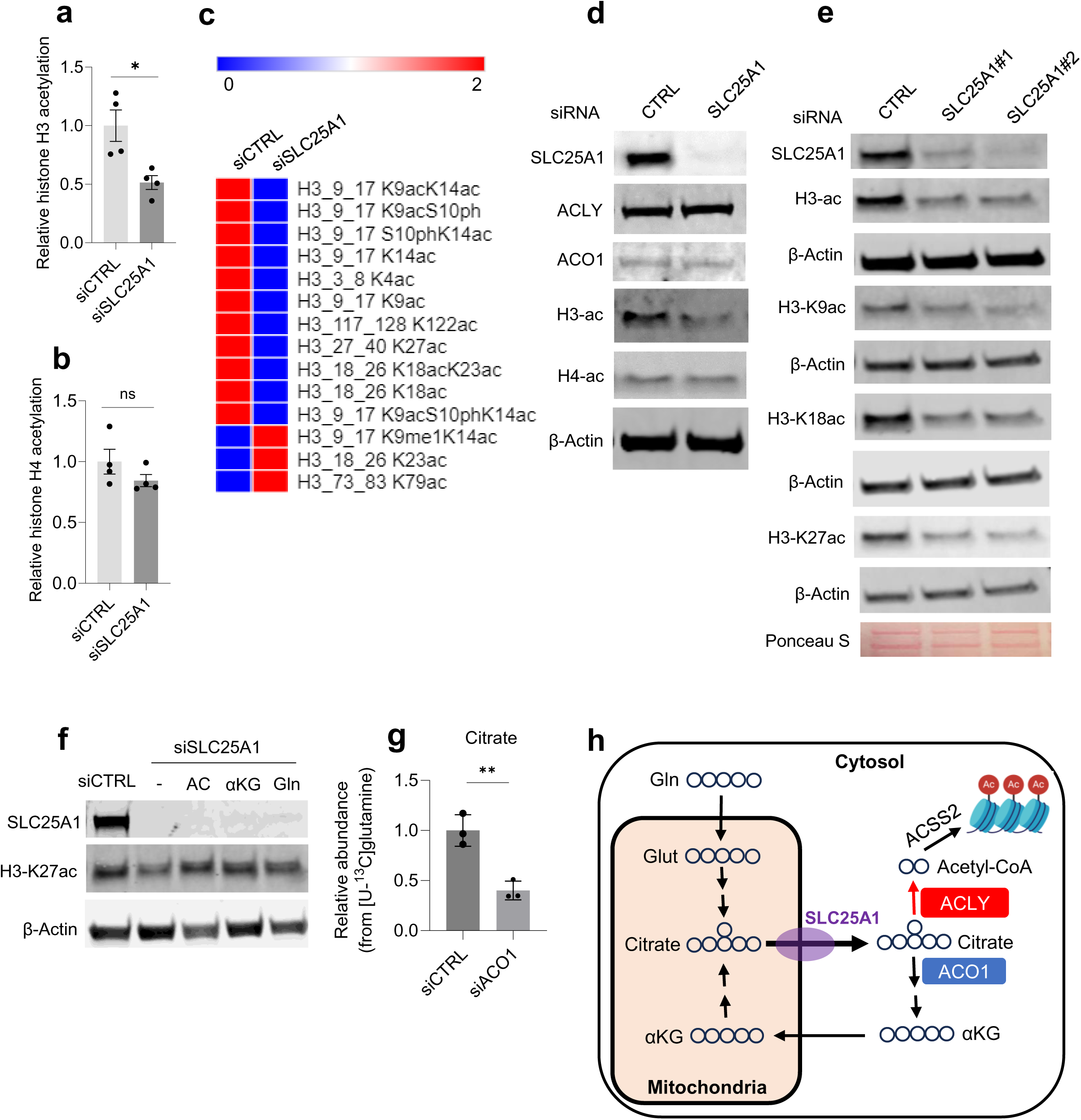
Citrate catabolism in cytosol maintains histone acetylation required for GPX4 expression under glucose limitation. Mass spectrometry-based measurements of total histone H3 acetylation (**a**), histone H4 acetylation (**b**), and heatmap showing differential modifications (**c**) associated with **a** in SLC25A1 knockdown cells cultured in medium containing 1 mM glucose for 30 hours. **d,** Immunoblots of SLC25A1, ACLY, ACO1, total histone H3 acetylation, total histone H4 acetylation, and control in SLC25A1 knockdown cells and control. **e,** Immunoblots in cells transfected with two independent siRNA against SLC25A1. **f,** Immunoblots of SLC25A1, H3-K27ac, and control. SLC25A1 knockdown cells were incubated with sodium acetate (AC, 4 mM), aKG (4 mM), or glutamine (Gln, 4 mM) for 48 hours. **h**, Total abundance of citrate pool from [U-^13^C]glutamine in ACO1 knockdown cells and control. **i,** Schematic diagram of glutamine-derived metabolites. Gln, glutamine; Glut, glutamate. n=3 independent biological replicates. Data provided as mean±s.d. Pairwise comparisons were conducted using two-tailed, unpaired Student’s *t*-tests. *representing *p*<0.05, **representing *p*<0.01.

To determine how αKG regulates histone H3 acetylation, we hypothesized that the entry of ACO1-mediated αKG production to the mitochondria sustains metabolic flux between cytosol and mitochondria, which in turn, maintains the pool of citrate required to support ACLY activity. Indeed, metabolic tracing using [U-^13^C]glutamine revealed that total citrate levels were reduced in ACO1 knockdown cells compared to the control (**Fig. 4g**), uncovering the role of ACO1 in maintaining the pool of citrate in cells under nutrient stress (**Fig. 4h**). These results were corroborated by evaluating the status of histone H3 acetylation in ACO1 knockdown cells, further validating the role of ACO1-dependent pathway as a regulator of histone acetylation in PDAC cells under glucose withdrawal (**Extended Data Fig. 3b**).

Given that αKG availability regulates dioxgenases, including TET (ten-eleven translocation) family of methylcytosine dioxygenases and JmjC domain-containing histone demethylases^29,30^, which influences methylation levels on DNA and histones, we sought to investigate whether upregulation of histone or DNA methylation, due to dysregulation of ACO1-dependent pathway, impacts GPX4 expression. Indeed, SLC25A1 knockdown induced an increase in DNA methylation as detected by elevated 5-methylcytosine (5-mC) (**Extended Data Fig. 3c**); however, 5-azacytidine was unable to rescue GPX4 expression (**Extended Data Fig. 3d**), and these results were further validated with mass spectrometry characterization of histone (**Extended Data Fig. 3e-h**). Together, these data suggest that the lack of histone acetylation is likely the underlying mechanism of reduced GPX4 expression in the context of SLC25A1 suppression under glucose limitation.

### SLC25A1-GPX4 axis supports mitochondrial function under metabolic stress

Given cytosolic-generated NADPH is unable to cross mitochondrial membrane^31–33^, mitochondria rely on their own compartmental antioxidant defense machinery to detoxify excess ROS, allowing them to operate normally. We next determined the contribution of SLC25A1 to maintaining redox in the mitochondria. A surge in mitochondrial-derived ROS levels was determined upon SLC25A1 suppression (**Fig. 5a**). Reduced SLC25A1 activity also impacted mitochondria more broadly, as both mitochondrial respiration and TCA cycle function were identified to be reduced. Importantly, the effects were rescued by supplementing cells with the products of SLC25A1-mediated pathways (**Fig. 5b, c**).

**Figure 5.**
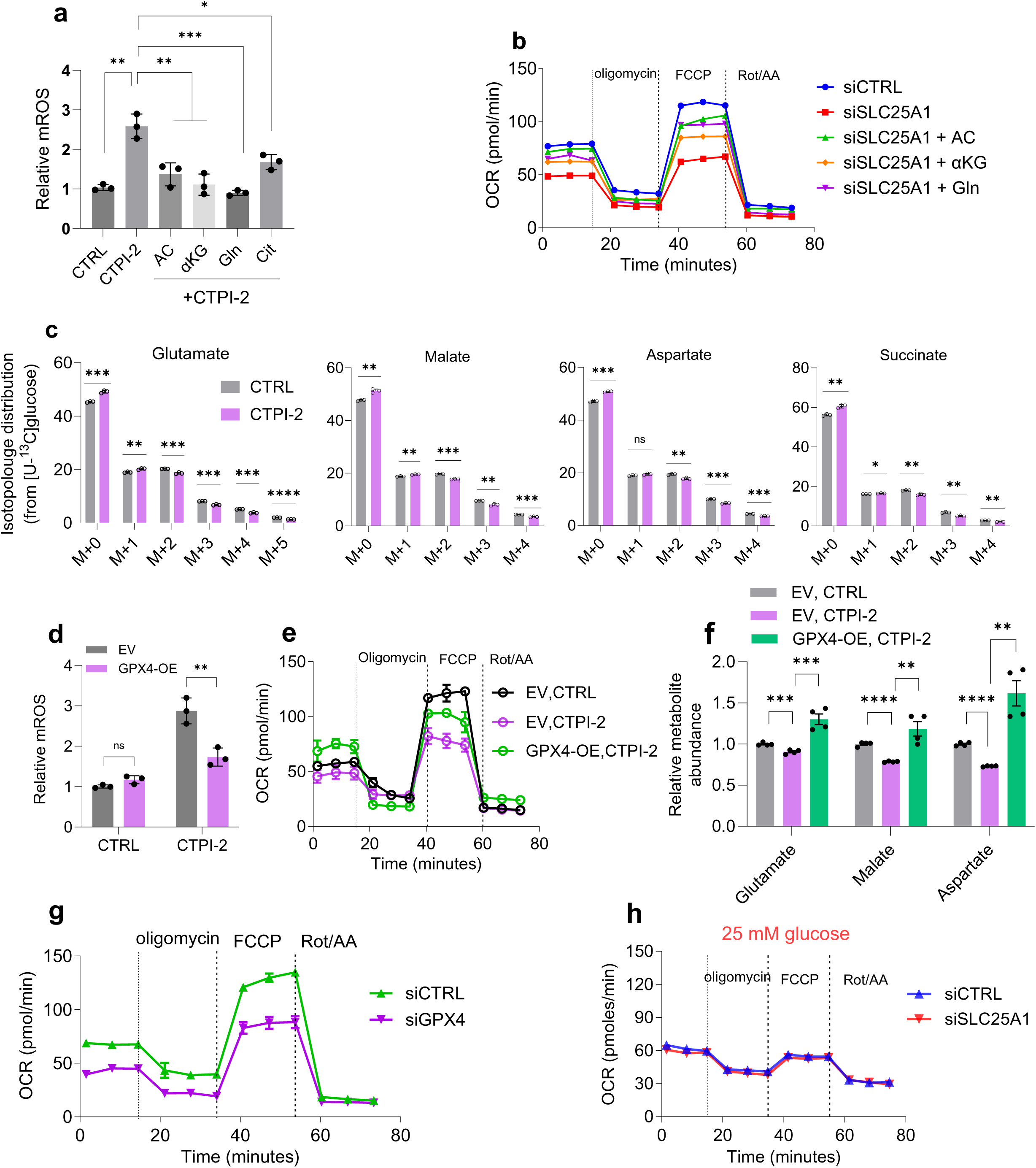
SLC25A1-mediated GPX4 expression supports mitochondrial function under nutrient stress. **a,** Relative mitochondrial-derived ROS measured with MitoSOX Red assay in cells treated with indicated compounds for 48 hours. **b**, Oxygen consumption (OCR) measurements in SLC25A1 knockdown cells and control incubated with sodium acetate (AC, 4 mM), α (4 mM), or glutamine (Gln, 4 mM) under glucose-restricted conditions (1 mM glucose) for 24-30 hours (representative experiments of two independent biological replicates with similar results are shown). C, Isotopologue distribution of metabolites derived from [U-^13^C]glucose in cells treated with CTPI-2 (100 µM) or vehicle for indicated hours. Relative mitochondrial-derived ROS (**d**) and OCR levels (**e,** representative experiments of two independent biological replicates with similar results are shown) in cells transfected with empty vector or GPX4 overexpressing vector were subjected to treatment with CTPI-2 (100 µM) or vehicle for 48 hours or 30 hours, respectively. **f,** Steady-state levels of metabolites measured by GC-MS in indicated cells treated with CTPI-2 or control for 30 hours. **g,** OCR measurements in GPX4 knockdown cells (**e**) and parental MiaPaCa-2 cells transfected with empty vector or GPX4 overexpressing vector treated with CTPI-2 (100 µM) or vehicle for 48 hours (g) and in SLC25A1 knockdown cells under glucose abundance (25 mM) (**h**, representative experiments of two independent biological replicates with similar results are shown). n=3 independent biological replicates, unless indicated. OCR data are representative of two independent biological replicates with similar results. Data provided as mean±s.d. Pairwise comparisons were conducted using two-tailed, unpaired Student’s *t*-tests. *representing *p*<0.05, **representing *p*<0.01, ***representing *p*<0.001, ****representing *p*<0.0001.

To determine whether induction of GPX4 expression via SLC25A1 is a critical regulator of mitochondrial function under metabolic stress, we performed metabolic profiling of cells in the absence or presence of exogenous GPX4. Upregulated ROS levels in cells under SLC25A1 suppression were substantially reduced by GPX4 overexpression. The rescue of GPX4 expression was also restored mitochondrial oxygen consumption (OCR) and TCA cycle intermediates (**Fig. 5d-f**), and the results were further validated by silencing GPX4 in cells (**Fig. 5g**). Notably, SLC25A1 silencing had no effect on mitochondrial OCR under nutrient abundance (**Fig. 5h**). These results establish a role for SLC25A1-mediated GPX4 expression that exhibits a global impact on mitochondrial function under stress.

### Elevated glutamine metabolism under SLC25A1 suppression

To turn SLC25A1 into a therapeutic target in PDAC, we performed a drug screening study where SLC25A1 inhibitor CTPI-2 was combined with over 4,000 compounds with known activity. Glutaminase inhibitor CB-839 was identified as one of the top hits that potentiates the growth inhibitory effects of SLC25A1 inhibition (**Fig. 6a**), suggesting glutamine metabolism as a potential compensatory mechanism under SLC25A1 suppression. Transcriptomic analysis identified upregulation of genes involved in glutamine uptake (**Fig. 6b**), and the results were further supported with labeling studies that showed increased enrichment of glutamine upon SLC25A1 inhibition, although the results did not reach the level of significance (**Fig. 6c**). The results from RNA sequencing also suggest upregulation of genes associated with glutamine metabolism. To this end, we evaluated whether glutamine metabolism is upregulated in cells lacking the activity of SLC25A1. Tracing studies using [U-^13^C]glutamine showed elevated enrichment of glutamine-derived metabolites, further highlighting the role of glutamine as a rescue mechanism under SLC25A1 suppression.

**Figure 6.**
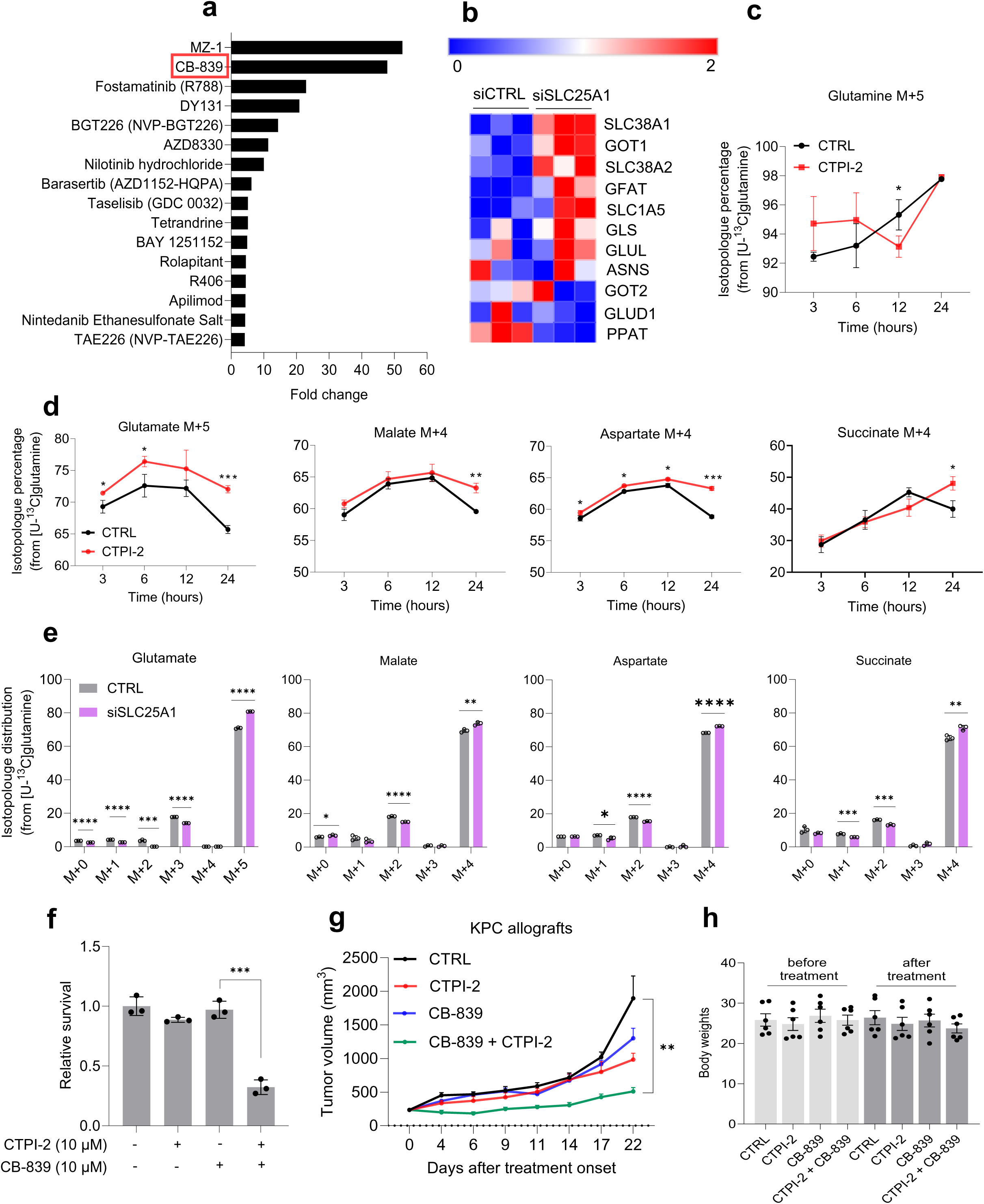
Enhanced glutamine metabolism upon SLC25A1 suppression. **a,** List of compounds, determined by drug screening, that increased sensitivity of MiaPaca-2 cells to SLC25A1 inhibitor CTPI-2 under low glucose conditions. **b,** Heatmap of genes associated with glutamine uptake and metabolism in SLC25A1 knockdown cells compared to control. Isotopologue distribution of glutamine M+5 (**c**) and indicated isotopolouges (**d**) derived from [U-^13^C]glutamine (4 mM) in cells treated with CTPI-2 (100 µM) or vehicle for indicated hours. **e,** Isotopologue distribution of indicated [U-^13^C]glutamine-derived metabolites in cells treated with CTPI-2 (100 µM) or vehicle for 24 hours. **f**, Relative survival of cells treated with indicated compounds for 5 days. Growth of subcutaneous KPC allograft tumors (**g**) and associated body weights of mice (**h**) under indicated treatments in a syngeneic mouse model of PDAC (n=5 mice per group; CTPI-2, 30 mg/kg, IP, once daily; CB-839, 200 mg/kg, BID). n=3 independent biological replicates. Data provided as mean±s.d. Pairwise comparisons were conducted using two-tailed, unpaired Student’s *t*-tests. *representing *p*<0.05, **representing *p*<0.01, ***representing *p*<0.001, **** representing *p*<0.0001.

Enhanced incorporation of glutamine carbons into TCA intermediates was also observed with SLC25A1 knockdown (**Fig. 6e**), providing the rationale for evaluating a combinatorial treatment with pharmacologic inhibitors of SLC25A1 and glutamine metabolism, as characterized by drug screening. As expected, cotreatment of CTPI-2 and CB-839 blocked the survival of cultured PDAC cells and suppressed tumor growth *in vivo*, and the combination therapy proved to be safe in animals.

## Discussion

The data presented herein provide mechanistic insights into PDAC adaptation to cancer-associated stress. Our studies revealed a crucial role for mitochondria in maintaining redox balance under metabolic stress. Targeting mitochondria with either pharmacologic inhibitors of ETC or by suppression of mitochondrial citrate transporter impaired intracellular redox balance, and citrate supplementation rescued cells from oxidative insults. We identified PDAC cells are vulnerable to SLC25A1 suppression under nutrient limited condition, the feature of pancreatic TME. Providing cells with antioxidant NAC partially restored the phenotypes associated with the lack of SLC25A1 activity. These results are consistent with prior observations that SLC25A1 controls redox homeostasis under cancer-associated stresses^24,34^.

Nutrient limitation is a characteristic of pancreatic TME^1–4^. Understanding metabolic dependencies of cancer growth under harsh metabolic state can assist the development of an effective therapeutic approach in PDAC. Mechanistically, we showed mitochondrial-derived citrate through citrate transporter SLC25A1 provides substrate to both cytosolic ACLY- and ACO1-dependent reactions to induce GPX4 expression required for PDAC cell protection from ferroptosis when PDAC cells experience nutrient stress. Indeed, OXPHOS inhibitors dramatically suppressed GPX4 expression under nutrient stress in PDAC cells, uncovering a molecular mechanism for previously observed induction of ferroptosis upon OXPHOS inhibition in cancer cells^35^. The upregulation of GPX4 was identified at both mRNA and protein levels when PDAC cells were exposed to nutrient-restricted conditions. Our studies also revealed that mitochondrial citrate-induced GPX4 expression plays a crucial role beyond supporting redox homeostasis. GPX4 knockdown impaired mitochondrial respiration and metabolism, and partial rescue mitochondrial OCR with GPX4 overexpression indicates PDAC cell metabolism is reliant on GPX4 expression under glucose limitation, further establishing a crucial role for GPX4 as a metabolic dependency in nutrient-austere TME.

## Methods

### Chemicals

To inhibit mitochondrial citrate transporter, CTPI-2 (Cayman Chemical, no. 36933) was used. For rescue experiments, cell-permeable Citrate (Citric acid, Sigma, no. C2404), tri-Sodium citrate (Sigma, no. 1.11037.1000), Acetate (Sodium Acetate, Sigma, no. S2889), α (alpha-ketoglutaric acid, Sigma, no. K1128), L-glutamine (ThermoFisher Scientific, no. 25030081), NAC (N-Acetyl-L-cysteine, Sigma, no. A9165), Mito-Tempo (Sigma, no. SML0737), Fer-1 (Ferrostatin-1, Cayman Chemical, no. 17729), Z-VAC (Cayman Chemical, no. 14463), Trolox (Cayman Chemical, no. 10011659), Nec-2 (Necrostatin-2, Cayman Chemical, no. 11657), 3-MA (3-Mathyladenine, Cayman Chemical, no. 13242), and 5-azacytidine (Cayman Chemical, no. 11164) were used. For OXPHOS inhibition, Rot (Rotenone, Sigma, R8875) and AA (Antimycin A, Sigma, no. A8674) were used.

### Cell lines, cell culture, and reagents

All cell lines were obtained from ATCC, except murine PC cells (KPC K8484: Kras^G12D/+^; Trp53^R172H/+^; Pdx1-Cre)^1,8^. Mycoplasma screening was performed using a MycoAlert detection kit (Lonza). Cell lines were grown at 37 °C and 5% CO2. For standard cell culture, cells were grown in DMEM (25 mM glucose, 4 mM glutamine), supplemented with 10% FBS, 1% penicillin/streptomycin, and prophylactic doses of Plasmocin (Life Technologies, MPP-01-03). For experiments under low glucose conditions, glucose-free DMEM (Life Technologies, no. 11966) was titrated to the indicated glucose levels.

### Small RNA interference

Oligos were obtained from ThermoFisher Scientific with the following ID numbers: SLC25A1 (human, no. S13093, S13095), ACLY (human, no. S915, S916), ACO1 (human, no. S671, S672), SLC25A10 (human, no. S3661), SLC25A11 (human, no. 15945), SLC25A12 (human, 119659), SLC25A13 (human, no. S532445), SLC25A21 (human, no. S225961), and SLC25A22 (human, no. S36256). siRNA transfections were performed using Lipofectamine 2000 (ThermoFisher Scientific, no. 11668027).

### Quantitative RT-PCR

Total RNA was extracted using PureLink RNA isolation (Life Technologies, no. 12183025) and treated with DNase I (Life Technologies, no. AM2222) to eliminate DNA sources. cDNA was synthesized using 1 μg of total RNA, oligo-dT and MMLV HP reverse transcriptase (Applied Biosystems, no. 4387406). PCR reactions were performed in triplicate using ThermoFisher Scientific primers with the following ID numbers: SLC25A1 (human, no. Hs01105608_g1), SLC25A10 (human, Hs00201730_m1), SLC25A11 (human, no. Hs01087948_g1), SLC25A12 (human, no. Hs00186535_m1), SLC25A13 (human, no. Hs01573625_m1), SLC25A21 (human, no. Hs00229049_m1), and SLC25A22 (human, no. Hs00368705_m1). RT-qPCR acquisition was captured using a Bio-Rad CFX96 and analyzed using Bio-Rad CFX Manager 2.0 software.

### RNA-sequencing and analyses

RNA quality was assessed using an Agilent 2100 Bioanalyzer (Agilent Technologies). mRNA was purified from total RNA using poly-T oligo-attached magnetic beads. After fragmentation, the first strand of cDNA was synthesized using random hexamer primers, followed by the synthesis of the second strand of cDNA. The processes of end repair, A-tailing, adapter ligation, size selection, amplification, and purification were then performed. The resulting libraries were assessed for quantity using Qubit and real-time PCR, as well as for size distribution using a Bioanalyzer. The quantified libraries were pooled and sequenced on a NovaSeq X Plus (Illumina) sequencer using 150-bp paired-end format. FastQC was used to assess RNA-seq quality and TrimGalore was used for adapter and quality trimming. RNA-seq reads were mapped against hg38 using STAR (v2.7.9a) aligner with default parameters. DESeq2 analysis with an adjusted p value<0.05 was used to derive a list of differentially expressed genes. TPMCalculator was used to calculate TPM from BAM files.

### GPX4 overexpression

GPX4 overexpressing plasmids (Origene, human, no. SC208065) were used. Plasmid transfections were performed using Lipofectamine 2000 (ThermoFisher Scientific, no. 11668027). Transfection efficacy was evaluated with western blotting 48 hours post transfection in cells under complete medium with glucose abundance.

### Immunoblot analysis

Total protein was extracted with RIPA buffer (Pierce, no. 89900) supplemented with protease inhibitor (Life Technologies, no. 1861280) and quantified using the BCA Protein Assay (ThermoFisher Scientific). Proteins were loaded on Bolt 4-12% Bis-Tris Plus gels (Life Technologies, no. NW04120) and transferred to PVDF membranes. Membranes were probed with antibodies against SLC25A1 (Proteintech, no. 15235-1-AP), SLC7A11 (Proteintech, no. 26864-1-AP), ACLY (Proteintech, no. 15241-1-AP), ACO1 (Proteintech, no. 12406-1-AP), GPX4 (Proteintech, no. 67763-1-IG), and β-Actin (Proteintech, no. 66009-1-IG). Blots were probed with secondary antibodies customized for the Odyssey Imaging system Secondary antibodies (680RD Goat anti-Mouse IgG (Li-COR, no. 926-68070, 1:20000 dilution) and 800CW Goat anti-Rabbit IgG (Li-COR, no. 926-32211, 1:10000 dilution).

### Cell viability assays

Cell viability was estimated by MTT Cell Viability Assay (ThermoFisher Scientific, no. 15250061) or by crystal violet staining. For the latter assay, cell colonies were fixed by crystal violet dye upon completion of experiments. To determine relative growth, dye was dissolved in 10% acetic acid and the associated absorbance measured at 600 nm using a microplate reader.

### Intracellular ROS and mitochondrial-specific ROS measurement

To detect intracellular ROS and mitochondrial-specific ROS, cells were incubated with 10 µM H2-DCFDA (Invitrogen, no. D399) for 45 minutes and with 5 µM MitoSox Red (Invitrogen, no. M36008) for 30 minutes, respectively, in serum-free medium with relevant glucose level in a 96-well plate. The associated fluorescent intensity was measured a microplate reader.

### Lipid peroxidation measurement

To measure lipid ROS, a lipid peroxidation assay (Cayman Chemical, no. 10009055) was performed according to the manufacturer’s instructions. The associated absorbance measured at 600 nm using a microplate reader.

### Subcellular Fractionation

Nuclear and cytoplasmic extraction was performed using NE-PER nuclear and cytoplasmic extraction reagents (ThermoFisher Scientific, no. 78833).

### Histone isolation and mass spectrometry

Histones were purified using acid extraction, as previously described^38^. Briefly, adherent cells were cultured in 6-well plates and collected following treatment. The cell pellets were immediately resuspended in 0.2M H2SO4 and rotated at 4°C for 4 hours. After centrifugation, histones were precipitated from the supernatant by the addition of 100% w/v tricholoracetic acid (TCA) for at least 1 hour, followed by centrifugation. The pellet was washed once with acetone containing 0.1% HCl and finally with 100% acetone. Histone proteins were dried at room temperature. Total histones were subjected to chemical derivatization using propionic anhydride (Sigma-Aldrich) and digested with sequencing-grade trypsin at a 10:1 substrate:enzyme ratio for 6 hours at 37°C. The digested peptides were processed as previously described. A resolution of 60,000 was used in the Orbitrap for the full MS, followed by MS/MS spectra collected in the ion trap. Data were subsequently analyzed with in-house software.

### Oxygen Consumption Rates (OCR)

To measure mitochondrial respiration, cells were cultured in a complete growth medium in Seahorse XF HS miniplates (Agilent Technologies). Once completion of experiments, growth media was changed to XF assay medium and the plates were incubated at 37°C in a non-CO2 incubator for 45-60 min before starting OCR measurement using in-house Seahorse XF HS Mini (Agilent Technologies). The results were normalized to cell number.

### Metabolic Profiling

GC-MS analyses were performed using in-house Agilent 5977C Mass Selective Detector and an Agilent 8890 Gas Chromatograph equipped with a DB-5ms GC Column (30 m x 0.25 mm × 0.25 um, Agilent Technologies). Experiments were performed in biological triplicate. Cells were grown in complete growth medium in 6-well plates. Once cells reached 50% confluency, growth culture medium was changed to low-glucose medium for a 16-h incubation. Subsequently, for labeling studies, cells were incubated in medium containing either [U-^13^C]glucose (Cambridge Isotope Laboratories, no. CLM-1396-1) or [U-^13^C]glutamine (Cambridge Isotope Laboratories, no. CLM-1822-H-0.1) for 24 h. To evaluate the intracellular enrichment of citrate, cells were incubated in medium containing [1,5,6-^13^C_3_]Trisodium citrate (Cambridge Isotope Laboratories, no. CLM-10366-0). To extract polar metabolites, cell culture plates were washed with cold PBS three times on ice, and metabolites were extracted in 80% methanol using cell scrapper. Samples were evaporated with nitrogen gas at 37°C, and then, dried lysates were mixed with pyridine:MOX solution to protect keto- and aldehyde groups at 40°C for 1 hour. Next, metabolites were derivatized with TBDMS at 60°C for 1 hour, and subsequently, the samples were transferred into GC-MS vials for analysis. The column temperature was initially set at 70°C and held for one minute, then ramped 7.5°C/min until 250°C. Samples were then ramped 25°C/min until 300°C, and held for 15 min. Metabolomics data were analyzed using the MSD ChemStation Software, version: F.01.03.2365 (Agilent Technologies). Metabolite counts were normalized to L-Norvaline, as an internal standard, and to the cell number. Masses were monitored in scan acquisition mode.

### In vivo studies

All experiments involving mice were approved by the Loyola University Chicago Institutional Animal Care Regulations and Use Committee (protocol no. 2023-026). Mice were maintained on a 12-hour light/dark cycle at room temperature with 30%-50% humidity under pathogen-free conditions in the animal facility. Mice were received standard chow and nutrient-free bedding. eight-to-ten-week-old, C57 black mice (C57BL/J) were purchased from Jackson Laboratories. Both genders were used. For flank xenograft experiments, 5×10^5^ cells KPC K8484 were suspended in 200 µL of a PBS:matrigel solution (1:1) and injected subcutaneously into the right flank. Once tumors reached ∼150 mm^3^, mice were randomized for the treatments. Tumor volumes were measured twice per week using a caliper (volume = length x width^2^/2). Body weights were measured weekly. Based on our IACUC protocol, the maximal tumor burden is 1,500 mm^3^ and tumor volume in our animal was not exceeded.

### Statistical analysis

Data are provided as mean ± s.d. from three independent experiments. Unless stated otherwise, two-sided type I error levels of 0.05 are assumed for hypothesis tests. For pairwise *P* values were calculated using two-tailed unpaired Student’s *t*-tests using GraphPad Prism 9.

## Acknowledgments

We thank Nancy Zeleznik-Le for manuscript feedback and valuable support, and Katie Kozlowska for assistance with tumor xenograft experiments. This work was supported by Cardinal Bernardin Cancer Center Translational Research Seed Grant (A.V.-G.), Elsa U. Pardee Foundation (A.V.-G.), and Perritt Foundation (A.V.-G.).

## Contributions

**Conceptualization:** A.V.-G. **Investigation:** A.K., K.L., A.A., J.C., S.G., C.T., S.S., A.V.-G. **Resources:** S.S., J.M.S., A.V.-G. **Data curation:** A.K., A.A., J.C., S.G., C.T., S.S., A.V.-G. **Writing:** A.V.-G. **Writing – review & editing:** A.K., K.L., A.A., G.A., X.D., W.S., C.T., S.S., J.M.S., A.V.-G. **Funding acquisition:** A.V.-G. **Supervision:** A.V.-G.

## Data availability

The RNA sequencing data generated in this study have been deposited in the Gene Expression Omnibus (GEO) database under accession number GSE286514.

---

**Extended Data Figure 1.**
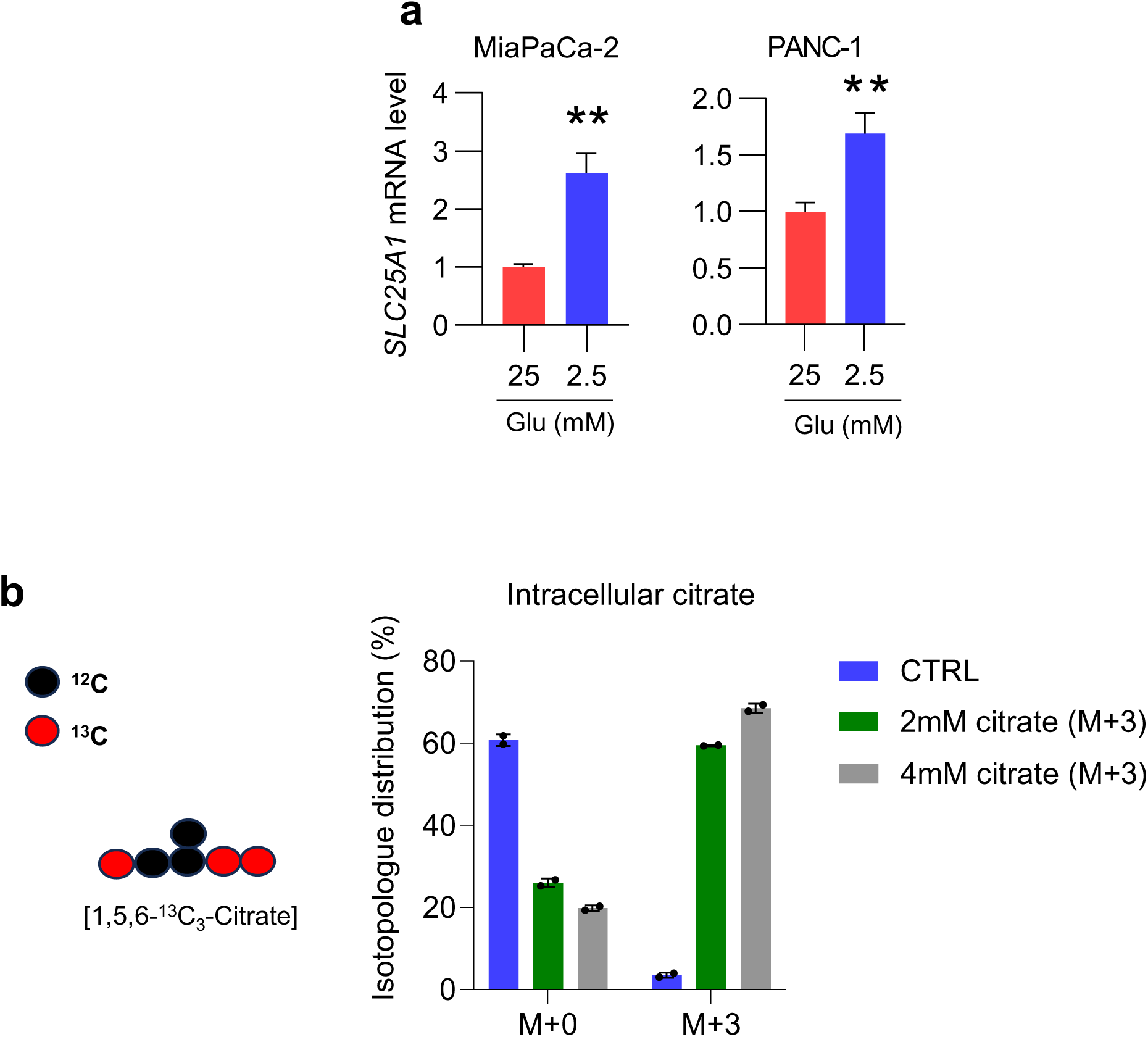
Increased intracellular enrichment of citrate after incubation for three hours. **a**, Relative mRNA levels of SLC25A1, normalized with 18S, in MiaPaCa-2 cells and PANC-1 cells cultured in medium containing either 25 mM glucose or 1 mM glucose for 24 hours for 48 hours. **b,** Distribution percentage of indicated isotopologues of citrate in cells incubated with [1,5,6-^13^C_3_-citrate] for three hours. n=2 independent biological replicates in **a**. **representing *p*<0.01.

**Extended Data Figure 2.**
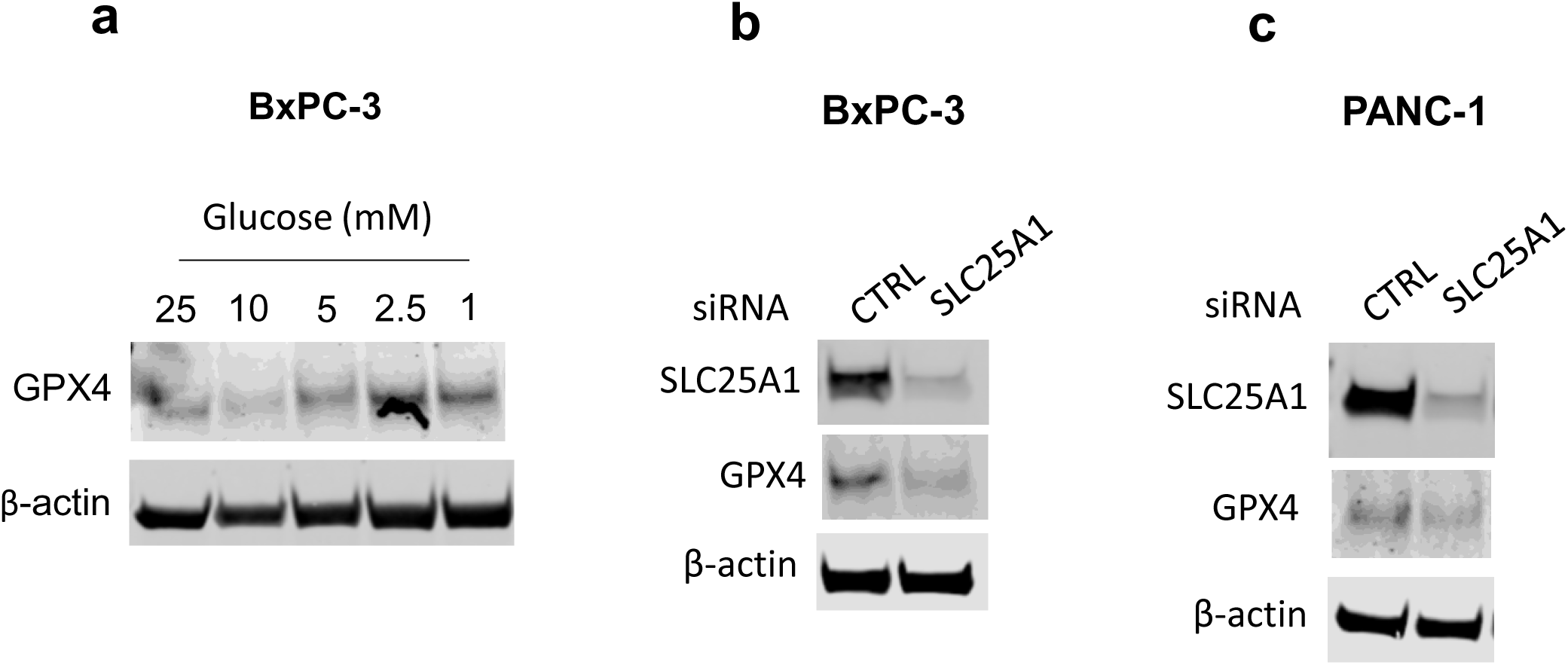
SLC25A1 and glucose availability regulate GPX4 expression in PDAC cells. Immunoblots of GPX4 and control in cells incubated with various levels of glucose for 48 hours (**a**), in SLC25A1 knockdown BxPC3 (**b**) and PANC-1 cells (**c**) compared to control. (n=2 independent biological replicates).

**Extended Data Figure 3.**
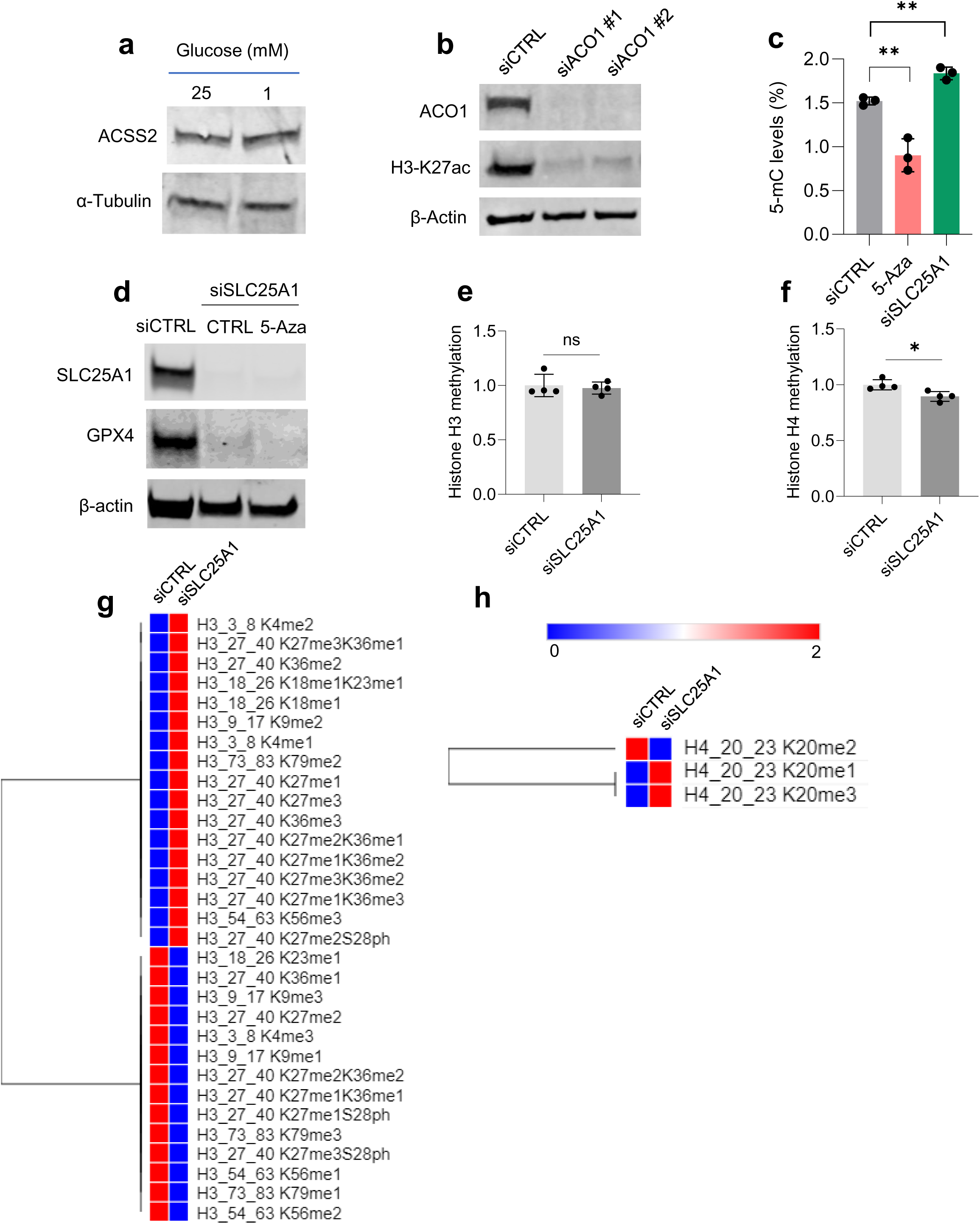
The impact of SLC25A1 knockdown on histone and DNA methylation. **a**, Immunoblots of ACSS2 and control in cells cultured in medium containing indicated glucose levels over 48 hours. **b,** Immunoblots of ACO1, histone H3 acetylation on lysine 27, and control in ACO-1 knockdown cells and control. **c,** Analysis of 5-methyl-cytosine levels percentage in SLC25A1 knockdown and control cells treated with 5-azacytosine (5 µM) cells cultured in medium containing 1 mM glucose for 24 hours. **d**, Immunoblots of SLC25A1, GPX4, and control in cells under indicated treatment for 48 hours. Mass spectrometry-based profiling of total histone H3 methylation (**e**), histone H4 methylation (**f**), and heatmaps showing associated modifications (**g, h**). n=3 independent biological replicates. Data provided as mean±s.d. Pairwise comparisons were conducted using two-tailed, unpaired Student’s *t*-tests. *representing *p*<0.05, **representing *p*<0.01

**Extended Data Figure 4.**
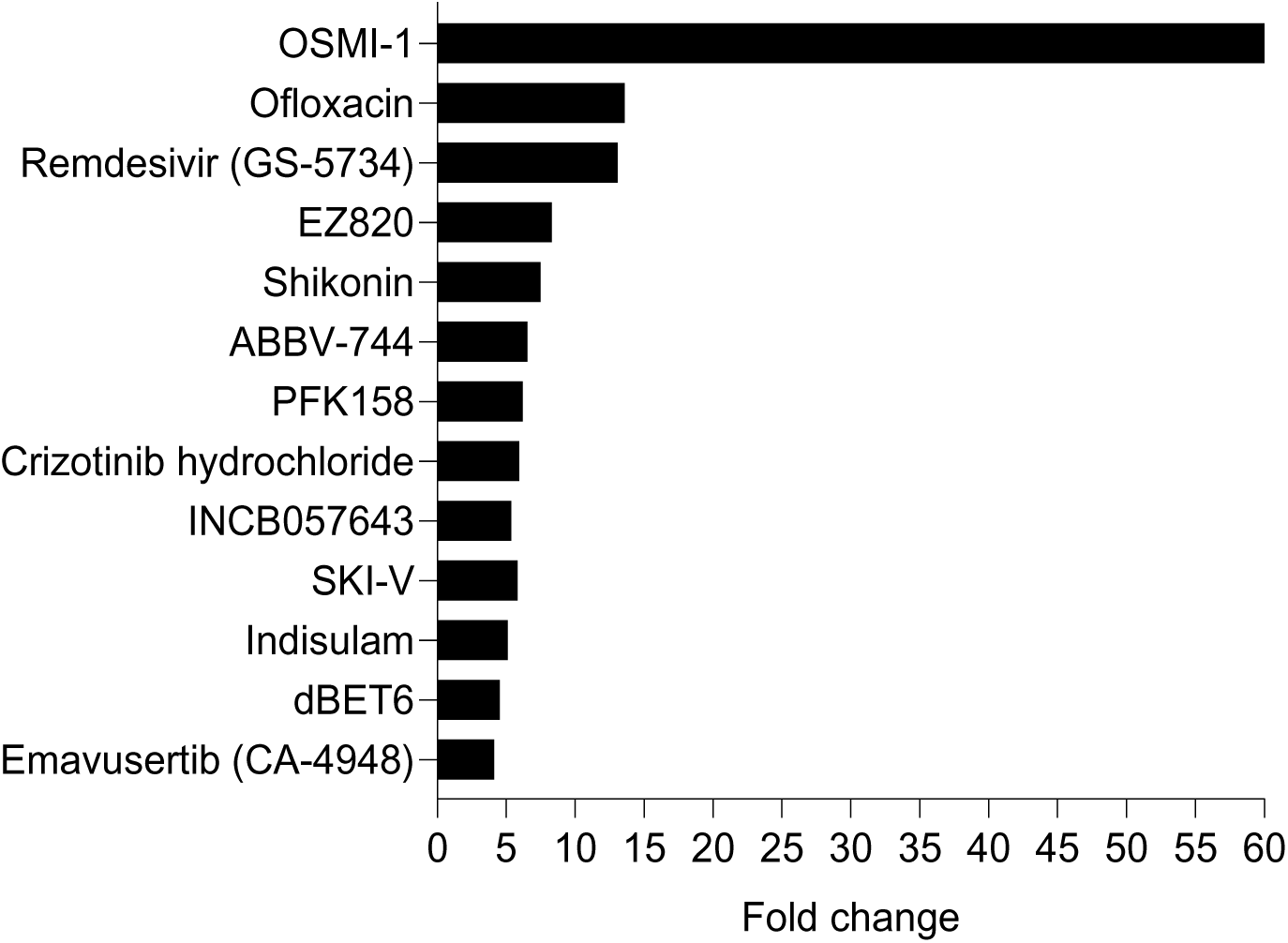
Distinct compounds sensitized PDAC cells to CTPI-2 under glucose abundance (25 mM glucose). List of compounds, determined by drug screening, that increased sensitivity of MiaPaca-2 cells to SLC25A1 inhibitor CTPI-2 under low glucose conditions.

